# Delay-dependent cholinergic modulation of visual short-term memory in rhesus macaques

**DOI:** 10.1101/2020.04.08.031377

**Authors:** Balázs Knakker, Vilmos Oláh, Attila Trunk, Balázs Lendvai, György Lévay, István Hernádi

## Abstract

Cholinergic neuromodulation is known to play a key role in visual working memory (VWM) – keeping relevant stimulus representations available for cognitive processes for short time periods up to a few minutes. Despite the growing body of evidence on how the neural and cognitive mechanisms of VWM dynamically change over retention time, there is mixed evidence available on cholinergic effects as a function of VWM delay period in non-human primates. Using the delayed matching to sample VWM task in rhesus macaques (N=6), we aimed to characterize VWM maintenance in terms of performance changes as a function of delay duration (across a wide range of delays from 1 to 76 s). Then, we studied how cholinergic neuromodulation influences VWM maintenance using the muscarinic receptor antagonist scopolamine administered alone as transient amnestic treatment, and in combination with two doses of the acetylcholinesterase inhibitor donepezil, a widely used Alzheimer’s medication probing for the reversal of scopolamine-induced impairments. Results indicate that scopolamine-induced impairments of VWM maintenance are delay-dependent and specifically affect the 15-33 s time range, suggesting that scopolamine worsens the normal decay of VWM with the passage of time. Donepezil partially rescued the observed scopolamine-induced impairments of VWM performance. These results provide strong behavioral evidence for the role of increased cholinergic tone and muscarinic neuromodulation in the maintenance of VWM beyond a few seconds, in line with our current knowledge on the role of muscarinic acetylcholine receptors in sustained neural activity during VWM delay periods.

## 1. Introduction

The importance of a visual stimulus is not necessarily commensurate with whether it is present in the current visual environment, or how much time has passed since it disappeared. To support adaptive behaviour, humans and animals have to maintain relevant stimulus representations available moment by moment despite the passage of time or changes in the environment. How different memory systems encode and maintain or recall events and factual information across various timescales is an important key question in current neuroscience. In the present study, we focused on visual working memory (VWM), the ability of retaining mental representations for short periods of time up to a few minutes due to its seminal function in general memory formation and its high clinical relevance, as decline in VWM performance is an early indicator of cognitive impairments in both healthy and pathological aging [1]. Ample research evidence on age-related cognitive decline has highlighted the role of cholinergic neuromodulation in VWM [2–4]. Among others, cholinergic neuromodulation has been and is still widely investigated as a promising symptomatic therapeutic treatment and is targeted for drug development, especially as part of currently evolving monotreatment regimes as well as combination therapies [5–7].

The delayed matching to sample test (DMTS) is a simple and highly translatable paradigm for the study of working memory that has been extensively used in both animal models [8–11] and human studies [12–15]. The basic notion of the DMTS task is to first memorize a sample stimulus and maintain the information on its identity across a delay period, then to identify the memorized stimulus among an array of non-matching stimuli. A wealth of behavioural pharmacology studies examined the connections between VWM and the cholinergic system in rodent [3,16–18] and non-human primate [19–27] preclinical models. In primates, for example, the anticholinergic agent scopolamine generally decreased delayed VWM memory performance in a delayed non-matching to sample task [27]. In a different study, nicotine has been shown to improve DMTS performance at medium-to-long delays individually adjusted to the animal’s performance between 0-60 s in middle-aged and senescent adult macaque monkeys [26]. Moreover, the acetylcholinesterase inhibitor donepezil (by prolonging the availability of acetylcholine for neuronal communication) has been found to exert a dose-dependent improvement on performance accuracy of macaque monkeys in the DMTS task [28–30]. The combined application of the acetylcholinesterase inhibitor physostigmine with the α2 adrenergic receptor agonist clonidine also promoted VWM performance [31,32].

It is well known that longer delay periods in the DMTS task ensue lower performance accuracy, which provides a simple way to measure the decay and dynamics of working memory over time [33,34]. Earlier, Bartus et al. [25] reported specific delay-related scopolamine effects in rhesus macaques performing in a modified delayed response paradigm using matrix illuminated food boxes, however, their investigation spanned delays ranging up to only 10 s in four discrete increments (0, 2.5, 5, 10 s delay). Penetar and McDonough [35] also found delay-dependent effects induced by the muscarinic acetylcholine receptor antagonist atropine in macaques in a simple three-colour visual DMTS task using delays ranging from 0 to 16 s. These observations were contradicted by Taffe et al. [27] and Glick et al. [36], who reported scopolamine effects on accuracy that were uniform in the 0-64 s delay range tested in discrete delay increments (0, 8, 16, 32 and 64 s in [27], and 0, 2, 8, 32 s in [36]) in VWM tasks using only two choice stimuli (one matching, one non-matching). Thus, previous studies using discrete unified delay lengths in preclinical non-human primate models did not conclude on pharmacological interaction effects with delay length, as most of them used tasks that were less demanding and also used a narrower delay range, disallowing the measurement of the full course of the delay-accuracy curve and its pharmacological modulation. While several other studies have taken into account the length of the delay period, they usually did so by initially titrating the delay duration to calibrate the preferred performance level of the individual animal instead of adhering to the absolute time values, resulting in a rather relative definition of short, medium and long delays within the full range of 2 to 180 s [28–30,33]. Individualized definition of delay latency categories can be very useful to optimize the sensitivity of the tests to the main effects of the pharmacological manipulation, however, it certainly precludes from drawing general conclusions about how the observed drug effects may interact with the temporal dynamics of memory encoding and maintenance.

Moreover, corresponding human studies suggest specific VWM [37] and auditory WM [38] encoding and/or maintenance related effects of systemic scopolamine administration with 4, 12 and 3 to 18 s delay period durations (respectively) over its weak sedating potential and non-specific effects on attention, leaving the field open for conclusive studies on the dynamics and pharmacology of VWM in preclinical translational research using an absolute rather than relative (calibrated to performance) time scale for the delay period.

Here, using the DMTS task in rhesus macaques, we aimed to study how cholinergic neuromodulation influences VWM encoding and maintenance across a wide range of delays (1 to 76 s) that were fixed across animal subjects to investigate the potential existence of specific delay-dependent amnestic effects of scopolamine in primates. By using identical delays for all animals and assuming that – despite the potentially differing performance levels – the temporal dynamics of memory encoding and maintenance would be similar among animals, we sought to pinpoint in what phases of the delay period the well-established cholinergic neuromodulations [39,40] would most potently affect VWM performance. To do so, we measured the effects of the anticholinergic amnestic agent scopolamine, administered alone or in combination with two doses of the acetylcholinesterase inhibitor donepezil to probe for the delay-dependent reversal of scopolamine effects.

We hypothesized that, as implied by earlier clinical research [37], both scopolamine-induced impairments and donepezil induced reversal effects would be stronger at longer delays, also sparing very short delays that might rely on the lower-level sensory components of VWM as previously observed in monkeys [41].

## 2. Methods

### 2.1. Subjects

Six 5-year-old male rhesus macaques (*Macaca mulatta*) were included in the study, weighing 4.62 ± 0.25 kg at the beginning of the experiments. Animals were kept on a mild food restriction with closely observing their recommended daily metabolizable energy intake requirement (determined as 107 kcal*BWkg^-1^*d^-1^) [42]. Animals were fed once per day, in the afternoons, following the daily testing session. Diet was standard nutritionally complete lab chow specifically designed for non-human primates (Altromin Spezialfutter GmbH, Lage, Germany) and was daily supplemented with fresh fruits and vegetables. Water was available *ad libitum*. In the home cage and testing rooms, temperature and relative humidity were maintained at 24 ± 1 °C and 55 ± 5%, respectively.

All procedures were conducted in the Grastyán Translational Research Center of the University of Pécs. The study was approved by the Local and National Ethical Committees on Animal Research and the Department of Animal Health and Food Control of the County Government Offices of the Ministry of Agriculture (BA02/2000-11/2012). Measures were taken to minimize pain and discomfort of the animals in accordance with the Directive 40/2013. (II.14.): ‘On animal experiments’ issued by the Government of Hungary, and the Directive 2010/63/EU ‘On the protection of animals used for scientific purposes’ issued by the European Parliament and the European Council.

### 2.2. Delayed matching to sample paradigm

Animals individually performed the DMTS task (Figure 1A) in one session per day in large cubical testing compartments equipped with a 15-inch touchscreen with LCD monitor. For stimulus presentation and data acquisition we used the CAmbridge Neuropsychological Test Automated Battery (CANTAB) specifically designed for non-human primates (Monkey CANTAB Cognitive Testing system with Intellistation and Whisker software, Campden Instruments, UK). The stimulus set (camcog_mdms0) comprised 430 individual rectangular monochrome abstract binary images consisting of 16×16 rectangular pixels (see Figure 1A for example stimuli). The background was black, and the colour of the images could vary across trials, but all images within a trial had the same colour. The stimuli were 8.5 cm wide and 7 cm in height on the screen. Each experimental session consisted of 120 trials and lasted for approximately 60 min. Conditioning and experimental (drug) sessions were performed in a computerized operant testing chamber equipped with a touch screen and dry pellet delivery apparatus. In the testing chamber, the animals were free to move and access the touch panel.

**Figure 1.**
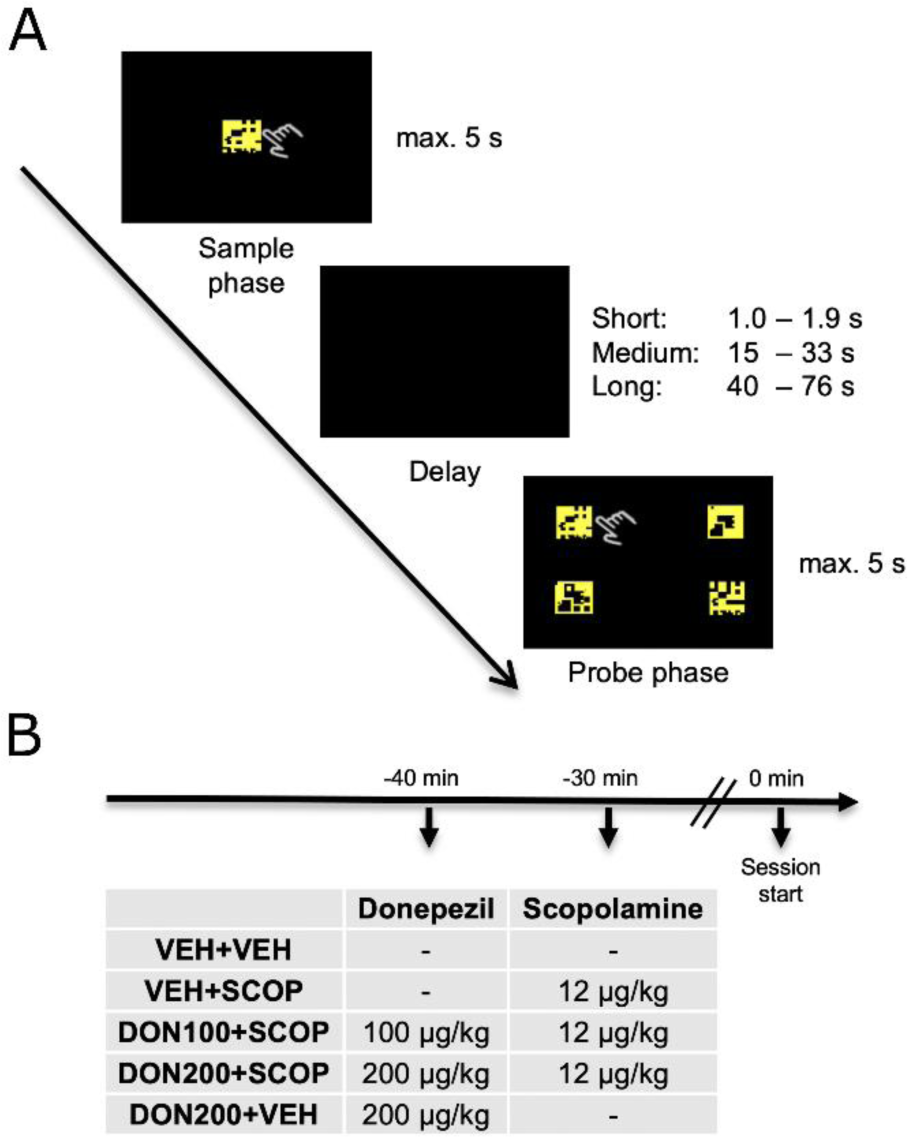
**(A) Schematic illustration of a single trial in the delayed matching to sample visual working memory paradigm and (B) combinations of treatments.** In the sample phase, the sample stimulus appeared in the centre of the screen, and the animals had to respond by touching the stimulus within 5 s. This was followed by a blank-screen delay period with 3 duration categories (short: 1 to 1.9 s; medium: 15 to 33 s; long: 40 to 76 s). After the delay had elapsed, 4 probe stimuli appeared in each quadrant of the screen, one of which was identical to the stimulus presented in the sample phase (target stimulus) and 3 were different (distractor stimuli). During this phase, the animals had to touch the target stimulus within 5 s. **(B) Schematic illustration of the treatment regime and timeline.** In each experimental session animals were administered acetylcholinesterase inhibitor donepezil (DON) and muscarinic receptor antagonist scopolamine (SCOP) alone or combined, or vehicle (VEH). Donepezil (or VEH) was injected at 40 min before the behavioural session, while SCOP (or VEH) was injected at 30 min before the session. We applied five treatment combinations as shown in the table.

Each trial consisted of a sample phase followed by a variable delay and a probe phase. At the beginning of each trial, a short tone was played to indicate the start of the trial. During the sample phase, a stimulus was presented at the centre of the screen that the animals had to touch. If the animal did not touch the sample stimulus within 5 seconds (sample response omission), the trial was considered unsuccessful and was terminated with no reward delivered. If the subject responded while the sample stimulus was still displayed, the sample phase was followed by a delay with blank screen. In the experimental sessions, we applied 3 delay duration categories: short delay between 1.0 and 1.9 s, medium delay between 15 and 33 s, and long delay between 40 and 76 s. There were 10 possible delay durations (delay bins) in each delay category: 1.0; 1.1; 1.2; 1.3; 1.4; 1.5; 1.6; 1.7; 1.8; 1.9 s for short delay, 15; 17; 19; 21; 23; 25; 27; 29; 31; 33 s for medium delay, and 40; 44; 48; 52; 56; 60; 64; 68; 72; 76 s for long delay. With 120 trials in each session, each exact delay duration was encountered 4 times, leading to 40 trials per delay category (short, medium and long) per session.

After the delay time had passed, the probe phase started, and 4 stimuli appeared in the four quadrants of the screen, one of which was identical to the stimulus presented in the preceding sample phase (target stimulus) and 3 were different (distractor stimuli). Upon touching the target stimulus within 5 s (correct response), the stimuli disappeared, and subjects received a pellet reward (190 mg Fruit Crunchies, Bio-Serv, Flemington, NJ) immediately after the response. In the case of an incorrect choice or if the animal did not respond within 5 s (probe response omission), the trial was considered unsuccessful and was terminated with no reward delivered. Each trial was followed by a 6±1 s inter-trial interval.

Before being recruited in the present study, all animals had been pre-trained to the final form of the task to performance criteria of at least 60% accuracy and over 80% motivation (trials completed) by having received excessive training in the DMTS task in over 100 experimental sessions with gradually increasing difficulty. After achieving the regular training criteria, the animals continued to routinely perform in the basic form of the task on every working (non-experimental) day with a constant 5 s delay duration. During the present experiments, all animals performed the task at the same time of the day on every weekday, including the treatment days.

### 2.3. Procedures and drug administration

In the placebo-controlled crossover and repeated measures experimental design applied in the present study, all subjects underwent 5 recording sessions covering all pharmacological treatment conditions with at least 72 hours of washout interval between the sessions. To achieve stable and high plasma levels at the time of task performance, intramuscular injections of donepezil (Gedeon Richter Plc., Budapest, Hungary) were administered 40 min prior to behavioural testing, followed by scopolamine (Tocris Bioscience, Bristol, UK) administered at 30 min before behavioural testing (Figure 1B). Donepezil and scopolamine were both dissolved in saline (0.9% NaCl); Saline was also used for vehicle (sham) treatments which were administered with the same timing as the genuine treatments (40 and 30 min before behavioural testing). Injection volume was set to 0.05 ml per kg body weight. All animals underwent five different treatments: vehicle + vehicle (VEH+VEH); vehicle + 12 μg/kg dose of scopolamine (VEH+SCOP); 100 μg/kg dose of donepezil + 12 μg/kg scopolamine (DON100+SCOP); 200 μg/kg dose of donepezil + 12 μg/kg dose of scopolamine (DON200+SCOP); 200 μg/kg dose of donepezil + vehicle (DON200+VEH). Each solution was freshly prepared before each recording session and was stored for less than two hours prior to administration. In initial scopolamine dose-finding pilot experiments, possible overt sedative side effects of scopolamine were subjectively observed by the experimenter. If the animal stopped responding during the task (which usually happened well before any overt sedative signs could be observed) we subsequently lowered the dose of scopolamine to preclude further overt sedative effects and to allow the desired continuous task performance. During the experiments, following a previously established rating scale [43,44] only normal or quiet arousal states were allowed (Scores 0 and 1) without drooping eyelids, slowed movements (Scores 2), intermittent sleeping (Scores 3), or complete sedation (Score 4). One animal showed higher sensitivity to scopolamine in terms of sedative side effects, so, to achieve similar amnestic effects (compared to the other subjects) without obvious side effects, a lower scopolamine dose (4 μg/kg) was used for him throughout the whole experiment.

### 2.4. Data analysis – repeated measures ANOVA

We analysed performance accuracy (number of correct responses divided by the number initiated trials – trials that had no sample phase response were excluded) and mean reaction time (time elapsed between the appearance of the probe stimuli and the animal’s response; RT). We analysed the data using two complementary approaches: conventional repeated measures ANOVA (rmANOVA) and Generalized Additive Models (GAMs).

In the rmANOVA analyses, data was pooled within each of the 3 delay bins (short, medium, long), and this 3-level Delay factor was used in all analyses to test for Delay effects and Treatment×Delay interactions. These analyses were conducted to foster comparability with previous studies, and to permit interpretation from a more traditional methodological perspective. First, a repeated measures factorial ANOVA was conducted to probe for scopolamine effects with the three-level Delay factor and a two-level Scopolamine treatment factor comparing VEH+VEH to VEH+SCOP. After establishing the scopolamine effects, further analyses were conducted to show where the DON+SCOP treatment conditions lie between the two extremes defined by VEH+SCOP (no reversal) and VEH+VEH (complete reversal), both on average (main effects) and also taking delay length into account (interactions). We were primarily interested in donepezil reversal effects, which were analysed in an rmANOVA with the three-level Delay factor and a three-level Donepezil treatment factor comparing DON100+SCOP and DON200+SCOP to VEH+SCOP; the null hypothesis here is that donepezil cannot counteract the effect of scopolamine. Afterwards, an additional analysis was conducted to test for residual scopolamine effects, using the three-level Delay factor and a three-level Donepezil treatment factor comparing DON100+SCOP and DON200+SCOP to VEH+VEH – the null hypothesis here is that donepezil can completely reverse scopolamine effects.

Control analyses were conducted on further dependent variables using the rmANOVA designs laid out above. Probe response omissions were counted as errors when calculating performance accuracy, so we analysed probe response rates (number of trials with responses in the probe phase divided by the number of trials with responses in the sample phase) to assess the contribution of response omissions to the main results. Sample phase response rates (number of trials with responses in the sample phase divided by the total number of trials) and mean sample phase RTs were also analysed as a proxy of potential lower-level effects on sustained attention, alertness or motor processes.

In rmANOVAs, Mauchly’s test showed that the sphericity assumption was not violated in any of the tests, so no correction was applied. Treatment×Delay interactions were further analysed using Holm’s method to correct to multiple comparisons. These analyses were conducted using the jamovi software [45].

### 2.5. Data analysis – Generalized Additive Models

Pooling the data into 3 large delay bins discards important information regarding the temporal dynamics of VWM performance that was the main focus of the present study. Simply using averages for each distinct value of delay length leads to only 4 trials per cell, therefore a method that effectively pools information across neighbouring delay values while also providing estimates with better temporal resolution is needed. Therefore, we also used Generalized Additive Models to estimate the delay-dependent pharmacological modulation of accuracy and RT in the VEH+VEH, VEH+SCOP, VEH+DON100 and VEH+DON200 treatment conditions.

In the GAM analysis, penalized thin-plate regression splines were fit using restricted maximum likelihood, as implemented in the `mgcv` R package [46,47]. For accuracy data, the dependent variable was an indicator variable (`Correct`) whether the given trial was performed correctly (1) or incorrectly (0), and the model was a binomial GAM with a logit link function. For RT data, a gaussian GAM was applied. Parametric main effects for Treatment were included in both models, also allowing treatment effects to vary within experimental subjects by adding a random term for Treatment. To model the interaction of pharmacological treatments with delay effects, the model was parametrized so that a smoothing spline was fit to a reference level, and for the remaining treatments difference smooths were fitted (analogously to intercept and simple treatment effects in a dummy coded linear regression). This is achieved in `mgcv` by specifying Treatment as an ordinal factor object, and using the following formula:

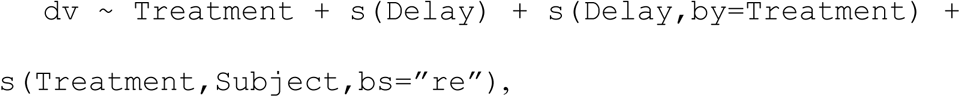

where `dv` is the dependent variable (`Correct` or `RT`), `Treatment` is the main effect term for pharmacological treatments, `s(Delay)` is the smooth term for accuracy in the reference condition as a function of delay length (as a standardized variable), `s(Delay,by=Treatment)` is the term for the differential smooths for the remaining levels and `s(Treatment,Subject,bs=“re”)` is the random effects term. To obtain comparisons of delay-accuracy and delay-RT functions (smooth interactions) for all pairwise comparisons of the 4 main pharmacological treatment conditions, we fitted 4 models, each using one treatment as reference level. Visual inspection of fits and model evaluation showed that the models using VEH+VEH, VEH+DON100 and VEH+DON200 as reference level yielded concordant results, while the model using VEH+SCOP as reference level provided different estimates and was also worse than the concordant ones in terms of Akaike’s Information Criterion (model with VEH+SCOP as reference: AIC_c_=3053, models with VEH+VEH, VEH+DON100, VEH+DON200 as reference level: AIC_c_=3051, small-sample corrected AIC calculated using the `AICc` function in the `MuMIn` package). Therefore, this latter model was not used in further analyses. All visualisations included are generated using the model with VEH+VEH as the reference treatment.

For each of the GAM models (3 for accuracy and 3 for RT), p values for Treatment effects and Treatment×Delay interactions were obtained using a permutation testing strategy as follows. From models fit on the observed data, statistics for the main effects (z values for accuracy and t values for RT, for each simple effect contrast) and the smooth interactions (χ^2^ for accuracy and F values for RT) extracted. Then, 999 surrogate datasets were generated by randomly shuffling the treatment condition label within each subject (but not across subjects). The above fitting procedure was then performed from the beginning on each of the surrogate datasets, yielding 999 of surrogate statistic values for each of the effects. These values were used as an empirical null distribution, based on the null assumption that our treatments are exchangeable with respect to the pharmacological effects and delay interactions that we observe. Specifically, permutation p values are derived by calculating the proportion of the permutation statistics showing an effect at least as strong as the observed effect, as detailed in [48,49]. Note that another possible strategy would have been to derive null distributions for each pairwise comparison, but in that case there would have been only 64 permutations, so we decided to prefer the more precise p-value estimates provided by a larger permutation distribution permitted by the currently applied less specific but still reasonable null hypothesis. In the cases when one comparison was performed with either of the compared treatments as reference level, we report the average of the two p values from the two models.

## 3. Results

### 3.1. Effects of delay duration in vehicle treatment

In the VEH+VEH condition, memory performance accuracy decreased continuously across the whole delay period, from 88% for small delay to 53% in the case of long delay (Figure 2; main effect of delay on accuracy: F_2,10_=16.84; p=0.0006, η_p_^2^=0.77; short vs. medium delay p_Holm_=0.032, medium vs. long delay: p_Holm_=0.016). Responses also got slower with longer delays (Figure 3; main effect of Delay on RT: F_2,10_=18.9; p=0.0004, η_p_^2^=0.79), dropping from 1800 ms to 2500 ms from small to medium delay (p_Holm_=0.0023) and decreasing at a slower pace to an average of 2700 ms in the long delay condition (medium vs. long: p_Holm_=0.20).

**Figure 2.**
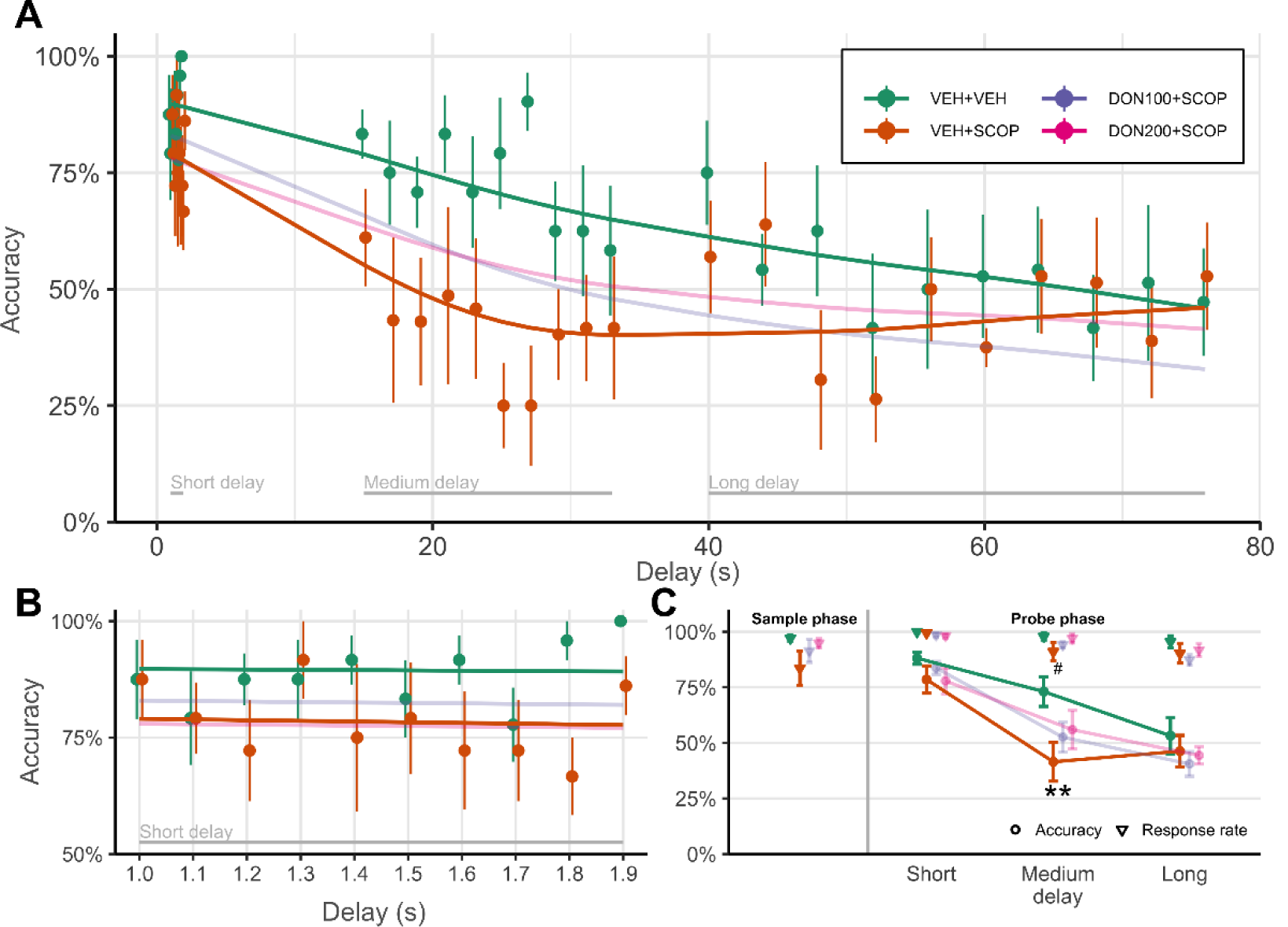
Delay-dependent scopolamine effects on performance accuracy in the DMTS task. **A:** DMTS performance as a function of delay in the VEH+VEH and the VEH+SCOP treatment conditions. Continuous curves indicate group-level estimates of accuracy from GAM fits. Markers and error bars show mean and s.e.m. across individual accuracies in the 30 small delay bins. The large delay categories (short, medium, long) are marked at the bottom of the figure. **B:** Accuracy estimates by GAM and small-bin averaging depicted focusing the short delay, with the same conventions as in *A.* **C:** Group-level average accuracies (circles) and response rates (triangles) in the sample phase and in the probe phase after short, medium or long delay. **: significant scopolamine effects for accuracy in relative to VEH+VEH in the medium delay, p_Holm_=0.0012; #: significant scopolamine effects relative to VEH+VEH for probe response rates, p=0.025 (rmANOVA main effect).

**Figure 3.**
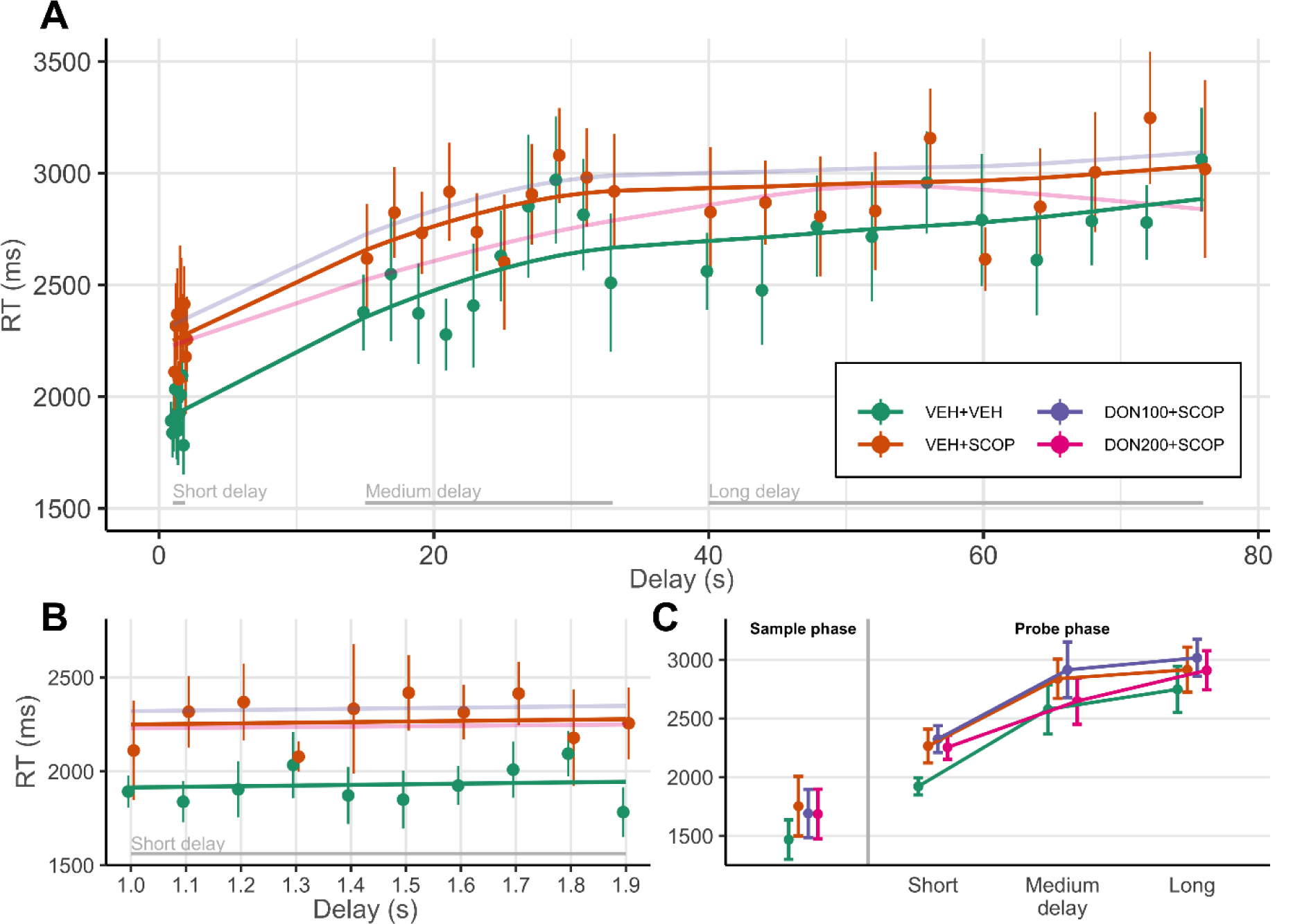
Scopolamine increased reaction time (RT) independently of task phase or delay length. **A:** Probe phase RT as a function of delay in the VEH+VEH and the VEH+SCOP treatment conditions. Continuous curves indicate group-level estimates of RT from GAM fits. Markers and error bars show mean and s.e.m. across individual RTs in the 30 small delay bins. The large delay categories (short, medium, long) are marked at the bottom of the figure. **B:** RT estimates by GAM and small-bin averaging depicted focusing the short delay, with the same conventions as in *A.* **C:** Group-level average RTs in the sample phase and in the probe phase after short, medium or long delay.

### 3.2. Scopolamine amplified delay effects on DMTS task performance

Scopolamine pre-treatment substantially deteriorated task performance on average (GAM main effect, VEH+VEH vs VEH+SCOP: p_perm_=0.001; rmANOVA main effect: F_1,5_=18.133, p=0.008, η_p_^2^=0.78) while also significantly changing the delay-accuracy function so that it reached its lowest point already in the medium delay period and did not decrease further (Figure 2; {VEH+VEH vs VEH+SCOP}×Delay smooth interaction: p_perm_=0.015; rmANOVA interaction: F_2,10_=5.213, p=0.028, η_p_^2^=0.51). Post hoc comparisons of the time-binned average accuracies in the VEH+SCOP condition from the rmANOVA also confirmed this (short vs. medium delay: mean ± s.e.m. difference: −37 ± 6.5% p_Holm_=0.0002; medium vs. long delay: −5 ± 6.5%, p_Holm_=0.85). As a result of this pattern, scopolamine effects were the strongest in the medium delay period (rmANOVA post hoc tests, VEH+VEH vs. VEH+SCOP, medium delay: −31.5 ± 6.1%, p_Holm_=0.0012) and were negligible when the delay period was short or long (VEH+VEH vs. VEH+SCOP, short delay, −9.6 ± 6.1%, p_Holm_=0.55; in short delay: −6.9 ± 6.1%, p_Holm_=0.82).

As a result of scopolamine treatment, RT increased by 250 ms on average (from 2360 ms to 2610 ms; see Figure 3) in a delay-independent manner, however, this effect was significant in the GAM analysis (GAM comparing RT in VEH+VEH with VEH+SCOP: p_perm_=0.0075, DON100+SCOP: p_perm_=0.0015, DON200+SCOP: p_perm_=0.038) but not in the rmANOVA (main effect of Scopolamine on RT: F_1,5_=3.68, p=0.11, η_p_^2^=0.42; Scopolamine×Delay: F_2,10_=0.22, p=0.80, η_p_^2^=0.04).

### 3.3. Donepezil partly reversed scopolamine-induced impairments in memory performance

Donepezil co-administered with scopolamine (DON100+SCOP and DON200+SCOP) partially reversed the impairments in accuracy caused by scopolamine (Figure 4). On average, donepezil treatment caused only a minor improvement in performance accuracy, yielding no significant differences relative to VEH+SCOP (GAM main effects, VEH+SCOP vs. DON100+SCOP and DON200+SCOP both p_perm_=0.24; rmANOVA comparing DON+SCOP accuracies with VEH+SCOP accuracy, main effect of treatment: F_2,10_=1.415, p=0.288, η_p_^2^=0.22). In addition, overall accuracy in DON+SCOP conditions was still substantially lower than the control accuracy levels (GAM main effects relative to VEH+VEH, DON100+SCOP: p_perm_=0.0035, DON200+SCOP: p_perm_=0.003; rmANOVA comparing DON+SCOP accuracies to VEH+VEH accuracy, main effect of treatment: F_2,10_=7.75, p=0.009, η_p_^2^=0.61).

**Figure 4.**
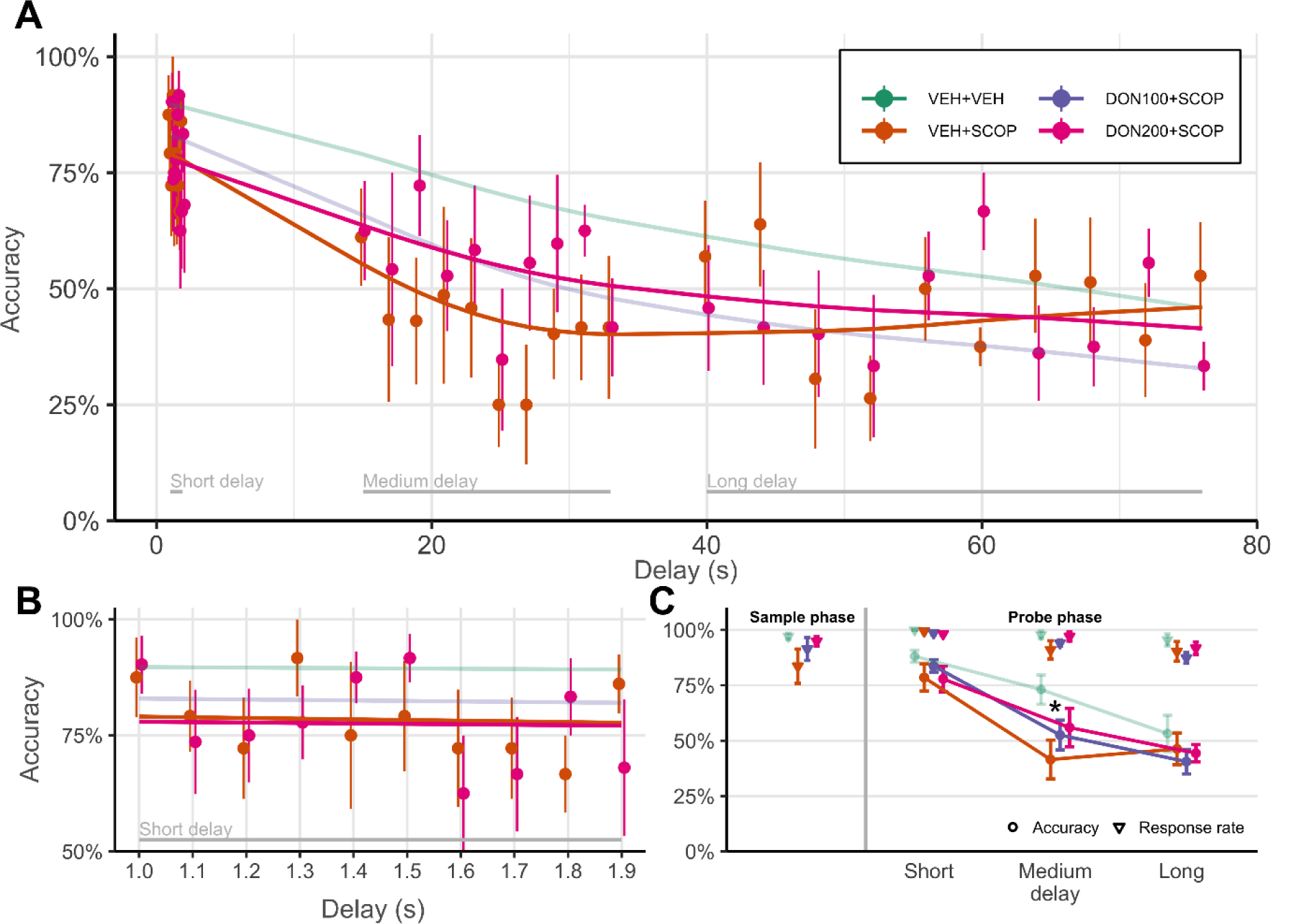
Donepezil partially reversed scopolamine-induced effects on performance accuracy. **A:** Delayed matching to sample task performance as a function of delay in the VEH+VEH and the VEH+DON200 treatment conditions. Continuous curves indicate group-level estimates of accuracy from GAM fits. Markers and error bars show mean and s.e.m. across individual accuracies in the 30 small delay bins. The large delay categories (short, medium, long) are marked at the bottom of the figure. **B:** Accuracy estimates by GAM and small-bin averaging depicted focusing the short delay, with the same conventions as in *A.* **C:** Group-level average accuracies (circles) and response rates (triangles) in the sample phase and in the probe phase after short, medium or long delay. *: p_Holm_=0.059

However, specifically for medium-length delays, performance accuracy was better than the corresponding VEH+SCOP performance by 11% for the 100 µg/kg donepezil dose (rmANOVA p_Holm_=0.33) and by 14% for the 200 µg/kg dose (rmANOVA p_Holm_=0.059; corresponding effects in the medium and long delay conditions ranged from 1% to 5%, p_Holm_=1). As a result of this small improvement, the shape of the delay-accuracy curve was largely reinstated, differing more from the VEH+SCOP shape (smooth interactions with Delay, VEH+SCOP vs. DON100+SCOP: p_perm_=0.042, VEH+SCOP vs. DON200+SCOP: p_perm_=0.088; rmANOVA Treatment×Delay: F_4,20_=2.985, p=0.044, η_p_^2^=0.37) and being relatively similar to the VEH+VEH shape (smooth interactions with Delay, VEH+VEH vs. DON100+SCOP: p_perm_=0.72, VEH+VEH vs. DON200+SCOP: p_perm_=0.234; rmANOVA Treatment×Delay: F_4,20_=1.74, p=0.18, η_p_^2^=0.258).

The DON+SCOP treatments did not show a dose-dependent difference neither in average accuracy (main effect, DON100+SCOP vs. DON200+SCOP: p_perm_=0.49; rmANOVA p_Holm_=1) nor in the shape of the delay-accuracy curve (smooth interaction, Delay×{DON100+SCOP vs. DON200+SCOP}: p_perm_=0.31).

An interesting additional finding to note is that in the GAM analysis DON200+SCOP responses appeared to be marginally faster than DON100+SCOP responses (p_perm_=0.085). This could be suggestive of a potential dose-dependent reversal effect, but other patterns of differences that would support this notion were absent, so we conclude that based on our data donepezil administered against scopolamine did not have an effect on RT.

### 3.4. Control analyses

Control analyses were performed on response rates and sample phase RTs. Probe response omissions were counted as errors in the main analysis, so it is important to assess the contribution of omissions to the results. While scopolamine decreased performance accuracy by 31% in trials with medium delays, the decrement in probe response rates was only 6% (see Figure 2C; in long and medium delay, on average, from 96% to 90%; main effect of Scopolamine: F_1,5_=9.9, p=0.025, η_p_^2^=0.67; Scopolamine×Delay interaction: F_2,10_=3.3, p=0.08, η_p_^2^=0.40). Donepezil tended to reverse the small scopolamine effect on probe response rates by a non-significant 3-6% improvement (see Figure 4C; Donepezil: F_1,5_=0.72, p=0.51, η_p_^2^=0.13; Donepezil×Delay: F_2,10_=1.36, p=0.28, η_p_^2^=0.21), which is nevertheless not comparable in magnitude to the 11-14% reversal effect on accuracy.

Trials without a sample phase response were excluded from the main analysis. However, to assess the motivational and alertness-related components in the observed pharmacological modulations, we conducted a separate analysis of sample phase response rate effects, as well. Analogously to probe response rates, scopolamine caused a 13.5% (non-significant) decrement in sample phase response rates (from 97.1% to 83.6%; see Figure 2C; main effect of Scopolamine: F_1,5_=2.8, p=0.15, η_p_^2^=0.36), which tended to be reversed by the (also non-significant) 8-11% relative improvement caused by donepezil (see Figure 4C; DON100: 91.4%, DON200: 94.9%; main effect of Donepezil: F_1,5_=1.71 p=0.23, η_p_^2^=0.25; Donepezil×Delay: F_2,10_=1.56, p=0.22, η_p_^2^=0.24).

Sample phase RTs were on average 1470 ms in the VEH-VEH control condition, and increased to 1760 ms in the SCOP+VEH condition (see Figure 4C; F_1,5_=5.04, p=0.075, η_p_^2^=0.50). These responses were similarly slow in both DON+SCOP conditions. We concluded that RT was unaffected by co-administered donepezil relative to VEH+SCOP (main effect of Donepezil: F_1,5_=0.30, p=0.75, η_p_^2^=0.06). This 290 ± 130 ms difference is comparable in both absolute magnitude and effect size (250 ms, η_p_^2^=0.42) to the probe phase RT scopolamine effect, and also with respect to the failure of donepezil to reverse this effect.

Donepezil monotreatment (DON200+VEH) compared to vehicle (VEH+VEH) did not show a significant effect of Treatment on accuracy (Treatment: F_1,5_=0.4, p=0.555, η_p_^2^=0.07; Treatment×Delay: F_2,10_=0.73, p=0.502, η_p_^2^=0.13). For RTs, while the main effect of treatment was not significant (F_1,5_=1.06, p=0.349, η_p_^2^=0.18), we found a significant interaction between Treatment and Delay (F_2,10_=7.152, p=0.012, η_p_^2^=0.59), which was probably due to a slightly stronger delay effect in the donepezil monotreatment condition, but no post hoc treatment effects were significant, and GAM analysis also did not support such conclusion.

## 4. Discussion

In the present study, we applied the DMTS VWM paradigm to investigate the effects of cholinergic receptor modulation on short-term memory of rhesus macaque monkeys in a delay-dependent manner. As also observed in the vehicle treatment condition in this study, continuous decrease in performance accuracy and increase in RT indicated that VWM representations deteriorated with the passage of the delay period. We investigated how the decay in accuracy of memory performance was influenced by cholinergic receptor agents administered before the testing session: we tested the transient amnestic effects of the muscarinic acetylcholine receptor antagonist scopolamine, and for the reversal of impairments, the acetylcholinesterase inhibitor donepezil was applied in combination with scopolamine.

We showed that scopolamine treatment decreased performance accuracy in the DMTS task, which reached an asymptotic low performance level earlier, with 20-30 s delay durations (medium delay category), while in the vehicle (control) treatment accuracy decreased continuously through the entire course of the delay period. This is in accordance with the pioneering results of Bartus et al [25], who observed dose-dependent scopolamine-induced deficits that were more pronounced with longer delay periods in a modified delayed manual response task. In their task, response accuracy in the control condition was very high (80-90%) even for the longest delay (10 s). In contrast, with delay periods up to 76 s and the more challenging task used in our study, we could measure the course of delay-dependent short-term memory deterioration even in the control condition, and also observe how this physiological VWM decay changed as a result of pharmacological treatments. Indeed, the considerably wider delay range (0-10 s vs, 1-76 s) used in our study revealed that under scopolamine treatment, performance reached the vehicle long-delay performance level already for medium-length delays, however, it did not decrease further. This is suggestive of a rightward instead of a downward shift of the performance accuracy curve, implying that it is primarily the pace, not the depth of delay-dependent short-term memory deterioration that changed as a result of scopolamine treatment.

Importantly – and also in line with previous results [25,50] – short-delay responses were only weakly or not affected by scopolamine. Short-delay performance is generally thought to reflect iconic memory [51,52] or more recently theorized intermediate memory states in this time range between iconic and *bona fide* working memory [53–57]. We suggest that these early VWM processes may be less reliant on muscarinic receptor-dependent mechanisms, in contrast to processes during or transitions into later states. The negligible effects for short-delay performance also excludes the possibility that scopolamine simply impairs visual discrimination of stimuli in the probe display [25].

Combined donepezil treatment was found to partially reverse the impairment caused by the administration of scopolamine with the same delay-dependent pattern, i.e. affecting performance accuracy only in the case of medium-length delay periods (15-33 s), partially reinstating the continuously decreasing performance pattern observed in the vehicle condition. There is abundant evidence that pro-cholinergic substances, including nicotinic agonists [58,59] and cholinesterase inhibitors [14,30,60] can mitigate scopolamine-induced deficits in short-term memory performance. Several studies by Buccafusco et al. [28,29,31–33] even used variable delay periods, however, in these studies, manipulating the length of the delay period was mainly used to titrate task difficulty for each animal to optimize sensitivity of the design to detect pharmacological effects, and not to explicitly characterize the delay-dependence of such effects. The resulting variability in delay length between animals in these studies exceeds the plausible range of between-subject variation in the course of cognitive events during VWM maintenance, and is rather likely to reflect generic, delay-independent factors behind each animal’s performance.

Previous studies have shown cognitive enhancing effects of donepezil monotreatment (without amnestic agent) in memory tasks in young and aged macaques [28,29]. Our present study does not confirm such enhancing effect: Here, we rather provide evidence for donepezil to reverse scopolamine-induced impairment of performance accuracy, and not for an additive enhancing effect.

Importantly, our control analyses on response rates support our interpretation that the cholinergic effects in this study reflect modulation of VWM processes. While probe response omissions (counted as errors in the main analysis) could hypothetically reflect both VWM-related and lower-level deficits, we showed that the contribution of omissions to the pharmacological effects on accuracy was minor. Sample phase response rates, where the role of memory processes can be ruled out, were also only weakly affected by pharmacological treatments. We suggest that the weak scopolamine effects on response rates that were almost completely reversed by donepezil reflect lower-level motor and alertness-related pharmacological modulations, including the well-known mild sedative-like effects of scopolamine. Taken together, control analyses make it clear that the main results on delay-dependent cholinergic effects on DMTS accuracy cannot be attributed solely to the lower-level effects on motor, motivational or alertness-related neural processes, but rather reflect genuine cognitive effects on the temporal dynamics of VWM.

Reaction times also depended on the length of the delay period, however, showed less sensitivity to pharmacological manipulations. Scopolamine induced a 250-300 ms response slowing that did not depend on the length of the delay, moreover, this slowing was even present in the sample phase responses. Also, this effect was not modified by co-administration of donepezil in any way. These results suggest that RTs as measured by our touchscreen DMTS task – in contrast to performance accuracy – mainly carry information on how generic motor and alertness-related processes are affected by scopolamine (as also shown in [61]). Reaction times are thought to mainly reflect the time required for visual search in the probe array, which is known to be faster for shorter delay since shorter-lived memory traces provide better search templates and attentional guidance than memories stored for longer periods of time [62]. Future experiments using instruments that are more optimal for precise RT measurements (here we used a touchscreen instrument and the animals could freely move within the testing compartment) may provide interesting further insights into how attentional guidance by visual working memory could be modified by cholinergic neuromodulation.

The most prominent model relates WM maintenance to sustained neural activity, which is confirmed by single unit [63,64] and non-invasive [65] measurements as well. Modelling [40] and experimental data [13,66,67] suggest that muscarinic receptor activity is essential for persistent delay activity during working memory, and it is known that scopolamine applied locally in the dorsolateral prefrontal cortex diminishes delay-period firing rate, memory-related activity patterns [68] and also delay-period activity as measured by fMRI [13] in humans. Based on this, we hypothesize that the cholinergic modulation of the temporal persistence of working memory by scopolamine and donepezil observed in this study is probably due to disruption and partial recovery, respectively, of the muscarinic mechanisms supporting the maintenance of persistent stimulus-coding delay activity in the frontoparietal memory networks.

Recently, behavioural and neurophysiological data has convergently shown that the substrate of working memory dynamically changes with the passing of time, shifting from primary sensory areas to networks of frontal and parietal associative cortical and subcortical areas [69–71]. This process is paralleled by traversing from detailed sensory images to more abstract, e.g. categorical representations [62]. Miller and Desimone [72] have shown that despite marked behavioural effects of scopolamine, neurons in the inferior temporal cortex, at the highest level of the visual representational hierarchy, remain unaffected by the drug. This is in accordance with our findings on the lack of cholinergic modulations early in the delay, when sensory cortical representations are thought to play a more prominent role in VWM maintenance. Relatedly, the dependence of mid-delay memory maintenance on muscarinic mechanisms is supported by the findings of Aggelopoulos and colleagues [73], who have shown that scopolamine specifically hinders the formation of categorical, more abstract stimulus representations that are more prominent later in the delay period and also pave the way towards long-term memory encoding [13].

In conclusion, in rhesus monkeys performing a DMTS VWM task, we have extended the classical results on delay-dependent cholinergic effects by using delay lengths up to 76 seconds. We showed that the delay-dependent scopolamine-induced impairments on VWM maintenance specifically affected the 15-33 s time range, suggesting that scopolamine increases (worsens) the normal deterioration of VWM with the passage of time. In addition, we also tested how donepezil, a clinically widely used cholinesterase inhibitor with high translational relevance, could partially rescue medium-delay scopolamine impairments. These results provide strong behavioural evidence for the role of increased cholinergic tone and muscarinic neuromodulation in the maintenance of VWM beyond a few seconds, in line with our current knowledge on the role of muscarinic acetylcholine receptors in sustained neural activity during VWM delay periods. Taking the length of the delay period into account can be a valuable component of basic and preclinical pharmacological research on the behavioural manifestations of delay-dependent cholinergic mechanisms and could deepen our understanding of short-term memory and its age-related impairments.

## Abbreviations

BW: body weight;
DMTS: Delayed Matching to Sample;
DON: donepezil;
GAM: Generalized Additive Model;
RT: reaction time;
SCOP: scopolamine;
VWM: visual working memory

## Acknowledgements

The authors would like to thank Judit Inkeller for valuable technical contribution. The authors also thank Anna Káldi, Annamária Traubert and Bence Petrovai for their assistance in animal care.

This work was supported by Gedeon Richter Plc. and the Hungarian National Brain Research Program of the National Research, Development and Innovation Office of the Hungarian Government (grant number ‘2017-1.2.1-NKP-2017-00002’) and the Hungarian Higher Education Programme for Excellence (Felsőoktatási Intézményi Kiválósági Program, FIKP) [20765-3/2018/FEKUTSTRAT]). The funding bodies did not influence the study design, the collection, analysis and interpretation of data and the decision to submit the article for publication.

## Author Contributions

V.O., A.T., B.L., G.L. and I.H. designed the research. V.O. conducted the experiments, V.O., A.T. and B.K. performed data analysis. V.O., B.K., B.L. and I.H. wrote and reviewed the manuscript.

## Data Accessibility Statement

The datasets generated during and/or analysed during the current study are available from the corresponding author on reasonable request.

